# The Speech Envelope Following Response in Normal and Hearing Impaired Listeners

**DOI:** 10.1101/2022.03.12.484064

**Authors:** Tijmen Wartenberg, Markus Garrett, Sarah Verhulst

## Abstract

The aim of this work was to investigate the perceptual relevance of the frequency following response to the syllable /da/ for speech intelligibility in noise based on age and hearing deficits. Recordings of the auditory evoked potential from young normal hearing (NH) and older individuals with both normal hearing and high-frequency (HF) hearing loss were analyzed. EFR metrics obtained in quiet and noise condition were calculated and correlated with speech reception. The envelope following responses were analyzed in terms of amplitude, latency and noise robustness. The response was first simulated to form predictions on the effect of cochlear synaptopathy and outer hair cell loss on the EFR. The experimental findings were in line with the computational predictions in the found observation that the EFR was reduced as a consequence of ageing and HF hearing loss. Both the audiogram and the speech EFR magnitude fell short in the individual prediction of SRT in stationary noise, but they accounted well for group performance. We also obtained within-group EFR latency with a cross covariance matrix. Validation of the method confirmed that speech EFR latency was predictive of click ABR Wave V peak latency. Moreover, statistical analysis not only showed that the robustness of the EFR obtained in the noise condition was dependent on the degree of high-frequency hearing loss in the older NH adults, but also dependent on the EFR magnitude in the NH younger adults. These findings provide evidence towards the important role of the EFR in speech-in-noise perception.

## Introduction

Auditory brainstem potentials provide researchers glimpses of sensory processing in the earliest stages of the auditory pathway. Studying these potentials is necessary to increase the understanding of the impacts of sensorineural deficits on speech perception, characterized by a degeneration of propagation of efferent sensory information to auditory cortices. The frequency following response (FFR) is used as case study to detect sensorineural deficits, because it reflects the sum of sustained phase-locked activity from subcortical sources to the periodicity in (complex) sounds (Young & Sachs, 1979). This phase-locked behavior represents a coding profile of synchronous instantaneous firing rates and slowly oscillating discharge rate of auditory nerve fibers and nuclei in the midbrain. Temporal fine structure (TFS) of sounds play a role in encoding the low frequency range of sounds of the fast component, below the phase locking limit of a few kHz, after which only the temporal envelope (TENV) remains encoded in the slower component (Carney, 2018; Joris & Verschooten, 2013). The FFR is phase locked to fine structure and slower temporal envelope modulations. One of these two features can be attenuated by either adding or subtracting responses to stimuli of different polarity, resulting in envelope frequency following responses (EFR) and spectral frequency following responses (TFS-FFR) (Aiken & Picton, 2008).

The EFR is important for suprathreshold envelope detection of narrowband or pure tone amplitude-modulated (AM) sounds and has therefore been proposed as a marker of cochlear synaptopathy, which is known to compromise detection of the temporal envelope through the loss of efferent connections between inner hair cells and auditory nerve fibers (Bharadwaj et al., 2014; Henry et al., 2014; Paul et al., 2016). A strong EFR relates well to a good performance in a variety of tasks, such as speech segregation, sound localization and pitch discrimination (Bharadwaj et al., 2015; Coffey et al., 2016), suggesting that a high synchronization power is beneficla for boosting speech intelligibility in adverse conditions. Several studies have also examined the role of EFR to syllabi or vowels (e.g., Easwar et al., 2015). The EFR to voiced speech contains information related to the fundamental frequency and higher harmonics, which is potentially important for vowel and speaker identification. Young listeners with an above average ability to perceive speech embedded in noise at normal conversation level at different ages, were found to have a stronger EFR and shorter delay than listeners with a below average ability (Anderson et al., 2010). There is some evidence that the effect of noise on the EFR is age-related (e.g., Ruggles et al., 2012), but quantification of this effect and evidence of an effect of NIHL, are still lacking. In the studies of Schoof & Rosen, (2016), speech EFR derived metrics were able to provide some clarity in addressing speech reception masked by a stationary and modulated noise maskers, but the pure tone audiogram proved overall more powerful than the EFR metrics in that study. The TFS-FFR was neglected in these studies, although it may contain perceptually relevant information, specifically at multiple higher harmonics. The TFS-FFR has been hypothesized to be play an important role in masking release, based on results from computer simulations and EEG experiments (Bidelman, 2016; Shamma & Lorenzi, 2013). There is evidence that the TFS-FFR to a vowel is associated with performance to SIN perception and is even a better predictor of speech reception in stationary noise than the EFR (Mai et al., 2018).

In recent work from our lab, the EFR to a rectangular stimulus was found to be a good predictor of HP speech reception in SWN in normal hearing listeners (Mepani et al., 2021). This stimulus was not only designed to boost synchrony to the stimulus envelope of midbrain nuclei to the stimulus envelope, but also to enhance differences based on cochlear synaptopathy, while being relatively insensitive to HF hearing loss. The aim here was to compare these findings for the ability to explain differences in speech perception performance by targeting the magnitude of speech FFR synchrony to the fundamental and higher harmonics. Starting from the point of simulations, predictions of speech perception based on the EFR and TFS-FFR were established with a biophysical model of the auditory periphery (Verhulst et al., 2018) to simulate the effects of auditory periphery deficits. EEG data from the dataset of was then processed to obtain comparative EFR and TFS-FFR waveforms. The EFR and TFS-FFR responses evoked by the syllable /da/ were analyzed for 44 subjects divided in three groups: young individuals with a normal audiogram, older normal hearing subjects with a normal audiogram and age matched individuals with mild to moderate sloped hearing loss, to isolate cochlear gain loss from ageing. Moreover, we developed a latency analysis and noise analysis to derive sensitivity of the EFR to noise, investigate the role of timing of the EFR and to demonstrate the subcortical origin of the speech EFR in this paper by comparing it to wave latencies of the click ABR.

If the understanding of FFR and potential perceptually relevant features in the FFR is improved, this may facilitate the development of individualized hearing aid algorithms for hearing restoration. Since the aim of these algorithms is to enhance the intelligibility of natural speech, it is in our interest to not only study the brainstem responses to semantically non meaningful envelope-modulated sounds designed in the lab, but also investigate the FFR to natural speech. Learning how the representation of the FFR is dependent on individual hearing pathologies is important for a better understanding as to why speech perception, especially in noise, is compromised and if it can be detected from the FFR.

## Methods

### Participants

Forty-four subjects gave their informed consent before participating in the studies, for which they were financially compensated. All participants were native German speakers. Fifteen young normal hearing (8f, age = 24.5 ± 2.3 years old), sixteen older normal hearing (8f, age = 64.3 ± 1.9 years old) and thirteen older hearing impaired (8f, age = 65.2 ± 1.8 years old) participated in the study. Subjects were not only selected based on age, but also on hearing impairments to account for high frequency sloped hearing loss. The division of the older participants into the two groups was based on the results of pure tone audiogram of the best ear. The pure tone audiogram was measured in a silent booth for both ears from frequencies ranging from 125 Hz until 10 kHz (FIG. 1) using Sennheiser HDA 200 headphones. Individuals with hearing thresholds of >20 dB HL at 4 kHz were categorized as (mild) sloped hearing impaired. Until 2 kHz, the audiogram profiles of all subjects were relatively flat and were checked for local dips. Also DPOAE measurements were performed in the subjects in complement to the audiogram. This categorized two yNH individuals as HI, 11 yHI individuals as HI, and all but one of the oHI group as HI, based on the estimated threshold criterion of 25 dB HL at 4 kHz.

**FIG. 1:**
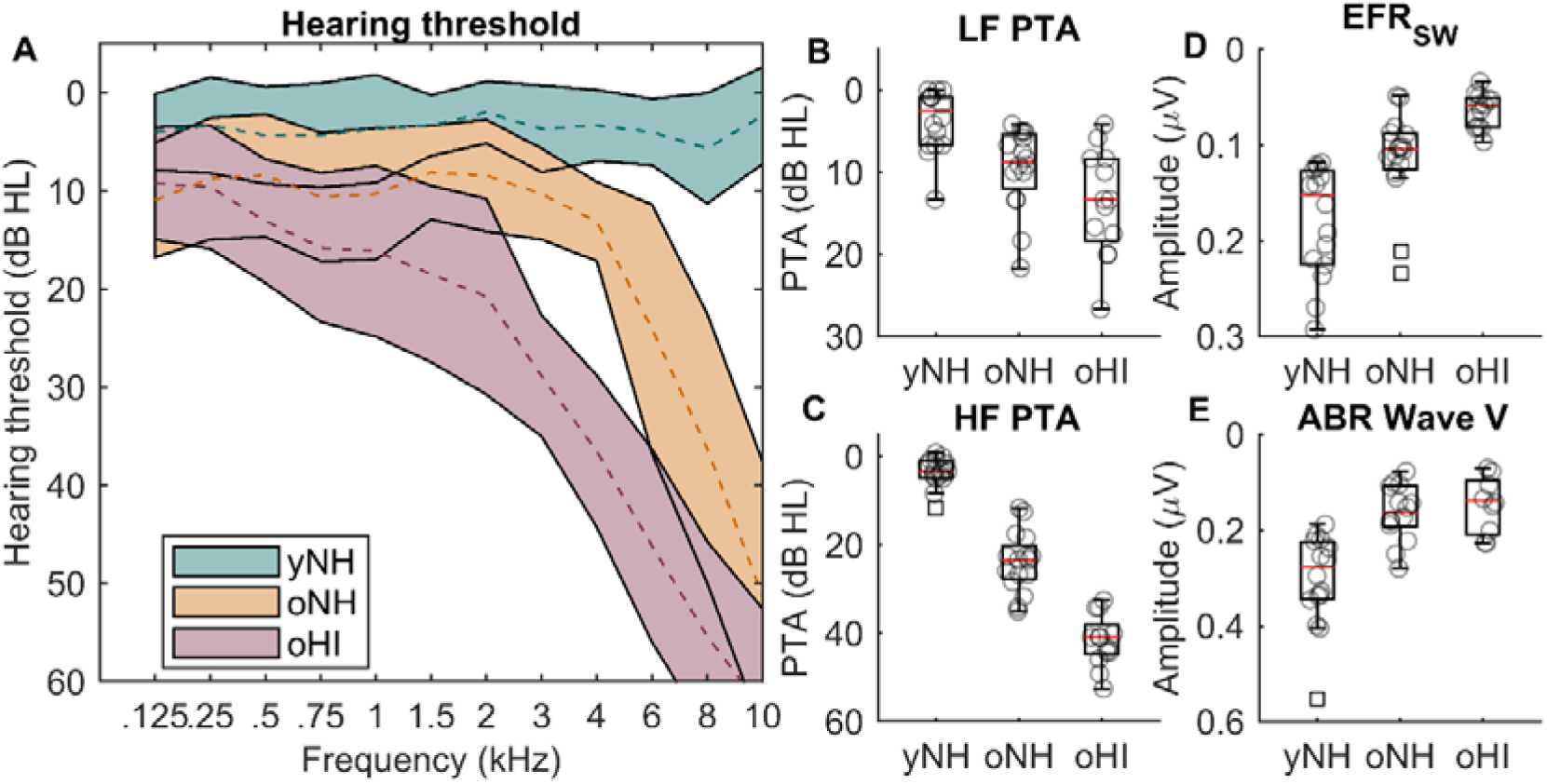
Audiogram : part of methods section. The audiometric thresholds of all subjects, divided over three groups, young normal hearing (yNH), older normal hearing (oNH) and older hearing impaired (oHI). Lines in A) denote average thresholds in different colors of the best ear for each group within the 95% confidence interval. The pure tone averages in B) and C) were formed by averaging the hearing thresholds in the regime of 125-1500 Hz for the lower frequencies and >1500 Hz for the high frequencies. The barplots in D) and E) illustrate differences in EFR strength to the square wave stimulus and the peak Wave V amplitude from the click ABR respectively.

**FIG. 2:**
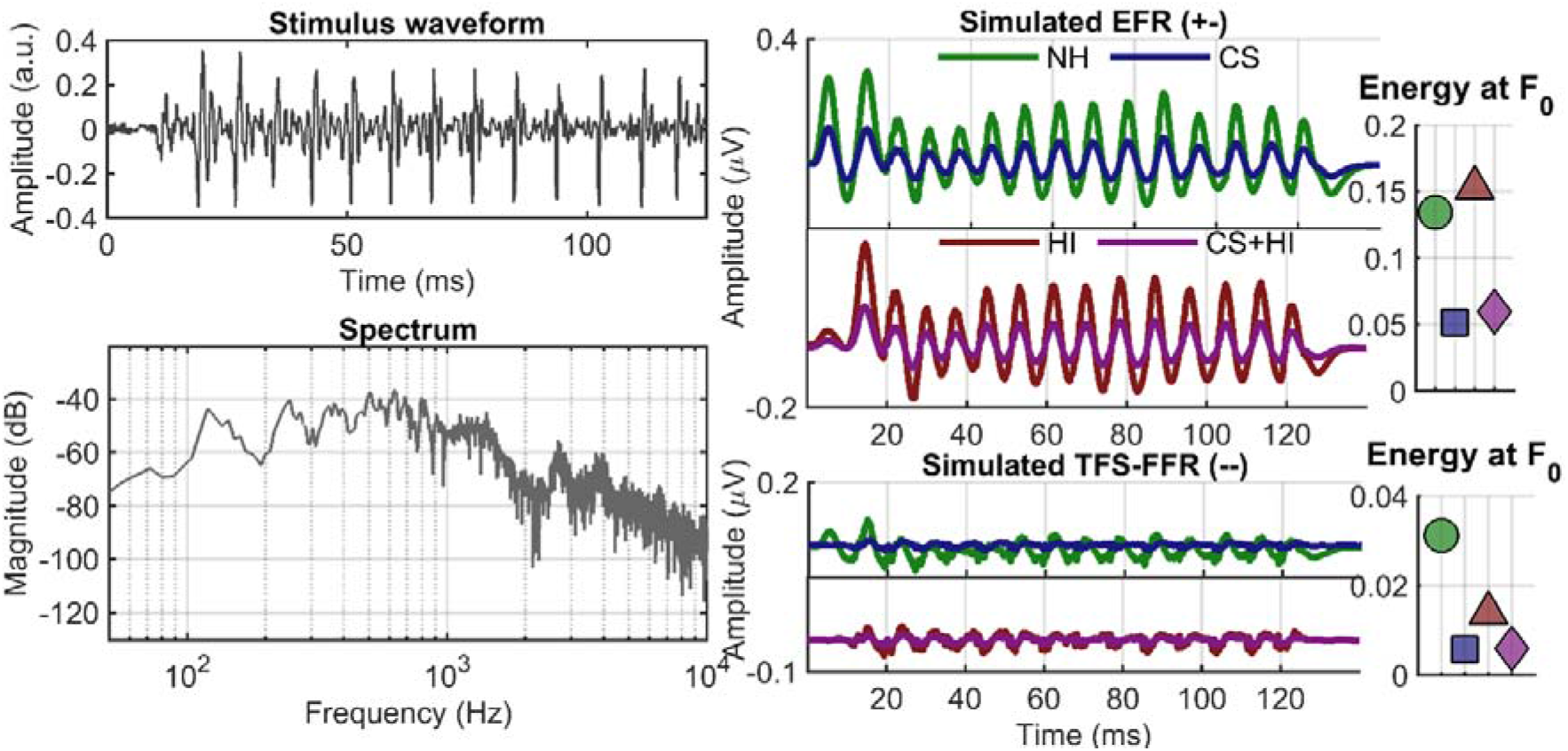
Simulations. Simulated time domain representations of the EFR (top) and TFS-FFR (bottom), modelled for the NH case (all fibers active and normal thresholds) and four cases of peripheral hearing deficits: CS (6 HS fibers), HI and a combination of CS and HI, with one the right side, an overview of the mean spectral energy at *F*_0_ in different symbols and in the same color scheme.

### Listening experiments

All subjects conducted a version of the German OLSA test (Kollmeier et al., 2015 for a review; Wagener et al., 1999) using the same headphones as for the audiometric measurements, but with monaural presentation in the best ear. Subjects were seated on a chair in a soundproof booth and independently performed a listening task on a laptop. During the task, as list was presented on the screen, which listed all possible choices of the presented five-word sentence, for example “Ulrich bekommt achtzehn grune schuhe” (*Ulrich receives eighteen green shoes*), giving ten options per word. For the speech in noise lists, the speech level was fixed at 70 dB SPL and the difficulty was adjusted to by step-wise varying the noise level after each decision in a one-up one-down criterion, based on the sentence score (Wardenga et al., 2015). The test was repeated for filtered sentences (FIR filter with order 1024) of the original sentences, by either low pass filtering the sentences at 1500 Hz or high pass filtering them at 1650 Hz (as in Verhulst & Warzybok, 2018). The comparison was also made more frequency specific by evaluating the speech reception threshold (SRT) in quiet and in noise for filtered sentences, allowing to segregate important high frequency cues from low frequency cues of sounds. The same tests were also performed in quiet. The signal level was then adapted to reach a performance of 50%. Performances in the matrix sentence tests are summarized in Table 1.

**TABLE 1:** Results of latency analysis. Results of the linear regression analysis to estimate click ABR W-V latency values, demeaned per group (yNH, oNH and oHI), with intergroup EFR latency for different combinations of subject groups.* : passes the statistical tests at the significance level of 0.05 **: passes statistical confidence after Bonferroni correction for six tests (k=6) performed per condition.

| Group | yNH | oNH | oHI | all | male | female |
| --- | --- | --- | --- | --- | --- | --- |
| Intercept (ms) | 0.25 | 0.21 | -0.47 | -0.12 | -0.48 | 0.84 |
| Coefficient | 1.20 | 0.95 | 0.98 | 1.09 | 1.18 | 1.24 |
| P | 0.0005** | 0.21 | 0.03* | <0.0001* | 0.01* | 0.004** |

### Computer simulations

The responses of the auditory periphery were simulated with a computational biophysical model (Verhulst et al., 2018b, v1.2.). A 100 kHz upsampled version of the syllable /da/ (FIG. 3.), same as in the experiment, was used as input to the model at a level of 70dB SPL for each polarity and several simulated hearing profiles, across 401 Greenwood spaced sections of the basilar membrane. The EFR was simulated by averaging synchronous responses from the auditory periphery opposite polarity stimuli, including contributions from three different ascending sources : auditory nerve fibers, cochear nuclei and inferior colliculi. The TFS-FFR was likewise simulated by subtraction. The NH profile was approximated by a weighted response from 13 HS, 3 MS and 3 LS fibers, and the CS profile was based on outputs from only 6 HS fibers to simulate the loss of IHC-AN ribbon synapses in cochlear synaptopathy. To simulate the sloped HI profile, 35 dB hearing loss at 8 k Hz starting at 1 kHz (see Verhulst et al., 2016)g, the poles of the BM admittance were adjusted to yield broader cochlear filters and reduce the nonlinear amplification of the simulated BM movement.

**FIG. 3:**
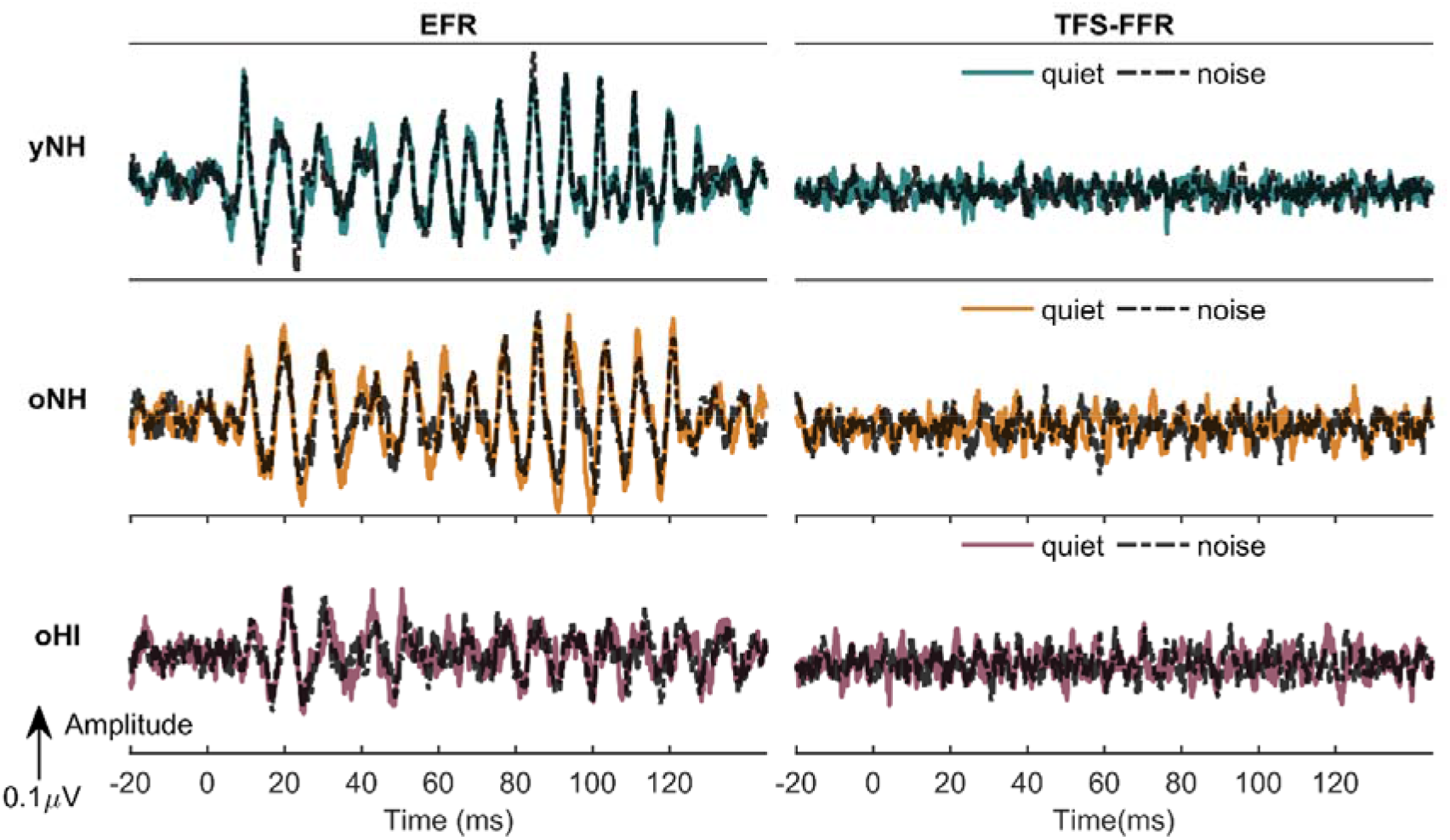
EFR & FFR. Grand average representations of the EFR and TFS-FFR for all three groups: younger normal hearing, older normal hearing and older hearing impaired shown on the same scales along the vertical axes and displayed from −20 ms relative to stimulus onset +20 ms relative to stimulus offset. Results from the recordings 35 dB SPL speech-weighted noise are plotted in darker lines on top of the results from the recordings in quiet.

### Protocol

#### Stimulus presentation

Sounds were delivered in the better ear through the magnetically shielded ER-2 insert earphone. The 120 ms-long syllable /da/ (F_O_ ~ 120 Hz), uttered by a male speaker, was presented 5000 times with a sampling frequency of 48 kHz at alternating polarity with an ISI of 100 ms. The stimulus was calibrated to be presented at a level of 70 dB SPL using an artificial ear simulator (type 4157, Bruel & Kjaer) . Experimental parameters were programmed in a custom MATLAB (R2015B) script. The computer for experimental control was connected to a FIREFACE UCX soundcard and a headphone driver (Tucker Davis), which was used in coherence with a custom EEG Trigger box by (University of Oldenburg) for synchronization between the EEG system and the sound delivery system. Electrooculography data was obtained with a 64-channel Biosemi system at a sampling frequency of 16,384 Hz.

The same set up was used to record responses to the rectangular stimulus at 70 peSPL and clicks at intensities of 70, 85 and 100 peSPL of alternating stimulus polarity. Specifics for these stimuli types in terms of design and analysis, are outlined in the study of (Mepani et al., 2021)

### Data analysis

#### Post processing

Differences in the current post processing protocol for analysis of the frequency following response of the speech stimulus to those of the rectangular wave, were in the processes of filtering of the responses, epoching and epoch rejection. All data was visually inspected and analyzed in MATLAB (R2015b). Data from channel *C_Z_* (frontocentral), referenced to the average electrode voltage across both earlobes, was zero phase filtered twice using digital FIR filters. The data was first highpass filtered at 80 Hz with a filter order of 1500, after which it was low pass filtered at 2500 Hz with a shallower filter order of 240. The cleaned data was consequently epoched from −20 ms relative to stimulus onset and +20 ms relative to stimulus offset. Epochs exceeding absolute values of >50nV were rejected from further analysis. In addition, epochs with total energy in the frequency region of interest (100-300 Hz) deviating more than two times from the median total energy across all trials, were also rejected from the analysis. The power spectral density was calculated with the Discrete Fourier Transform (DFT) and the total energy was summed within this region. Maximally 26% of all epochs was rejected for all participants with this procedure.

#### Envelope Following Response measures

The TFS-FFR and EFR were determined via the bootstrapping algorithm on the Discrete Fourier Transform of Zhu et al .(2013), combining information from both polarities, with the difference that the spectral magnitudes were not log-transformed to keep the amplitude in *μV*. Also, to calculate the TFS-FFR, the plus sign in the equation to add responses to opposite polarity, was switched to a minus sign to subtract the responses, following the argumentation in (Aiken & Picton, 2008). In the bootstrapping procedure, a total sampling size of 400 trials was used from both polarity trials, which was repeated 100 times. The noise floor calculation was then based on 200 repetitions. To finally calculate the TFS-FFR and TFS-EFR, the calculated spectral magnitude of the noise floor was subtracted from the raw spectral magnitude. The DFT calculation was performed after zero padding the signal to increase the length of the signals to 250 ms, resulting in a spectral resolution of 4 Hz. The energy at *F*_0_ and at multiples of *F*_0_ was calculated by averaging the energy within 8 Hz (two bins) around 120 Hz and multiples of 120 Hz.

Statistical validation was performed by transforming discrete Fourier SNR values relative to the spectral noise floor to z scores in the filtered region of 80-2500 Hz. All maximal values in the range 112-128 Hz were marginally higher than three for all subjects from the yNH group. Two z values were lower than 3 for the oNH group and four within the oHI group. The statistical confidence bound of z = 3 was reached for 20 subjects in total for the second harmonic. Z scores rarely exceeded this criterion in case of the TFS-FFR. Therefore, the TFS-FFR was rejected from further analysis. Also, since the energy at higher harmonics for the EFR are possibly the result of nonlinear interactions with no clear link to cochlear synaptopathy, only the energy at the lowest harmonic was analyzed when comparing the responses.

To achieve noise removal in the time domain, the post bootstrapped spectral magnitude was combined with the DFT phase angle of the pre bootstrapped averages to yield the new EFR and TFS-FFR estimates. In contrast to the method used in (Vasilkov et al., 2021), all bins were selected for the transformation to the time domain to keep an accurate representation of the slow temporal envelope. The operation was applied after bootstrapping was completed, instead of applying the operation on bootstrapped samples. The formulation for this processing on the FFR:

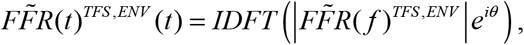

with *θ* the phase angle of the DFT of the original FFR.

#### Temporal alignment strategy

A simple cross-correlation between the response and the stimulus waveforms was found to not be conclusive enough about absolute EFR latency, so another simple but effective approach was used to temporally align the EFR waveforms of all individual waveforms to make a grand average representation of the EFR for the three group. The technique was designed to compensate for differences in delay of the onset of sustained EFR activity by temporally shifting the EFR of an individual and calculating the 3D cross-covariance matrix, including all other subjects within the group category in the range of 40 samples (τ ± 2.4 ms), for all n ∈ N individuals within each group category.

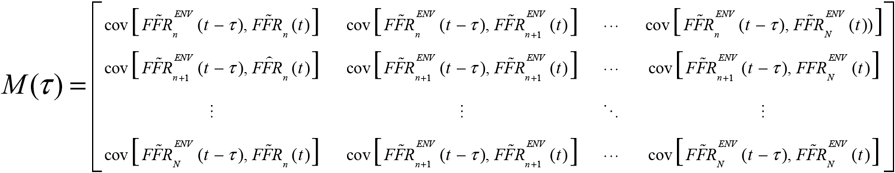

The optimal latency was determined as the lag corresponding to the maximal value in the average of the cross covariance matrix across all rows and for all time lags, τ:

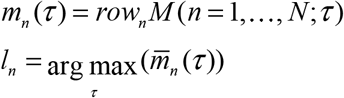

The defined lag window of 40 samples was wide enough to ensure a local maximum for all subjects, while preventing neighboring maxima roughly 136 samples (8.3 ms) apart, assuming a fundamental frequency of roughly 120 Hz. The utility of the latency values was to relate them to click ABR latencies for a source analysis of the speech EFR. Furthermore, the EFR waveforms were compensated for their latencies by temporally shifting them with −*l_n_* to sum all of the in-phase individual waveforms and as a result improve the visualization of the grand averages.

#### Noise robustness quantification strategy

Another measure was developed to estimate the robustness of the EFR to noise as the rms difference of the absolute error between EFR waveforms recorded in quiet and in background noise, after compensation of latency. The difference was normalized by the average rms of the EFR waveform. The noise ratio was defined as:

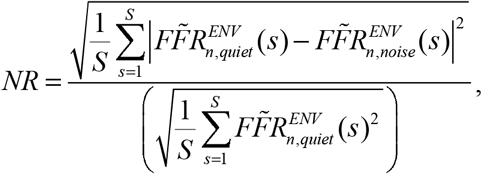

with s as sample (time in seconds / sampling frequency of 16384 Hz) and S the total length of the waveform in number of samples. Instead, the metric was designed to be mostly sensitive to asymmetry in local amplitudes during condensation and rarefaction periods of the wave-form and to minor asynchrony differences, or overall noisiness of the waveform. Thus, this measure was designed to be both a measure of robustness of the waveform to noise as well as robustness of the EFR across multiple recordings.

## Results

### Simulating frequency following responses

Brain potentials evoked by the speech stimulus in the experiment were modelled for four simulated different profiles to simulate the effect sensorineural hearing loss on the FFR (FIG. 4.) The EFR was reduced for the CS (cochlear synaptopathy) profile, which shaped the hypothesis that individuals in the oNH group had lower EFR amplitudes than individuals part of the yNH group. There was also a small elevation of the response for the HI profile, which suggested an additional small increase in EFR for the individuals in the oNH group in respect the oHI group. Based on the simulations, EFR magnitude was predicted to be a discriminant factor in for age, on the assumption that hidden hearing loss is common in the older population. The simulated TFS-FFR was found to be marginally higher for the NH profile, so size of the TFS-FFR was hypothesized to be a good indicator for HF sloping hearing loss.

**FIG. 4:**
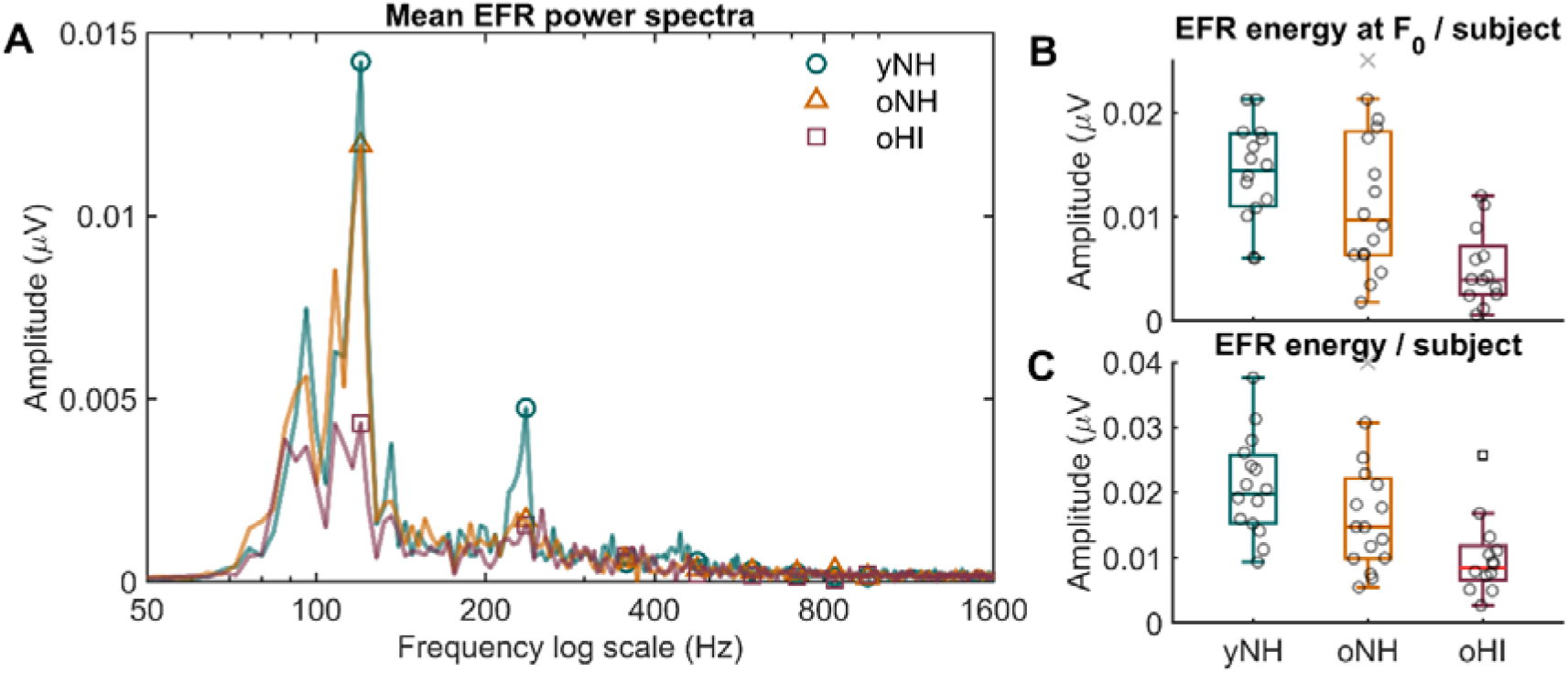
Box plots + energy of EFR/FFR. A:The mean power spectra of the bootstrapped EFR for the groups shown with a Discrete Fourier Transform with a spectral resolution of 4 Hz. Values at 120 Hz and harmonic are indicated with unique symbols for each subject category. B: Barplots of the energy at F_0_ for the three investigated groups. Individual results are denoted as circles. C: Same figures as B, but with the energy summed across the first five harmonics. This limit was based on visual inspection of the EFR power spectra.

### Derived frequency followed responses

Grand averages of the EFR and TFS-FFR for each group are shown in FIG. 5. In the representation of the EFR, there was baseline activity until the start of the presentation of the syllable, followed by a transient response after stimulus onset and rhythmic, sustained activity oscillating at the pitch frequency of the vowel /a/ (~120 Hz). The *EFR_oNH_* average retained much of the activity compared to the *EFR_yNH_* average, but the *EFR_oHI_* average was far more degraded. The TFS-FFR amplitude fluctuated around zero and showed little change in the post stimulus onset interval compared to baseline activity.

**FIG. 5:**
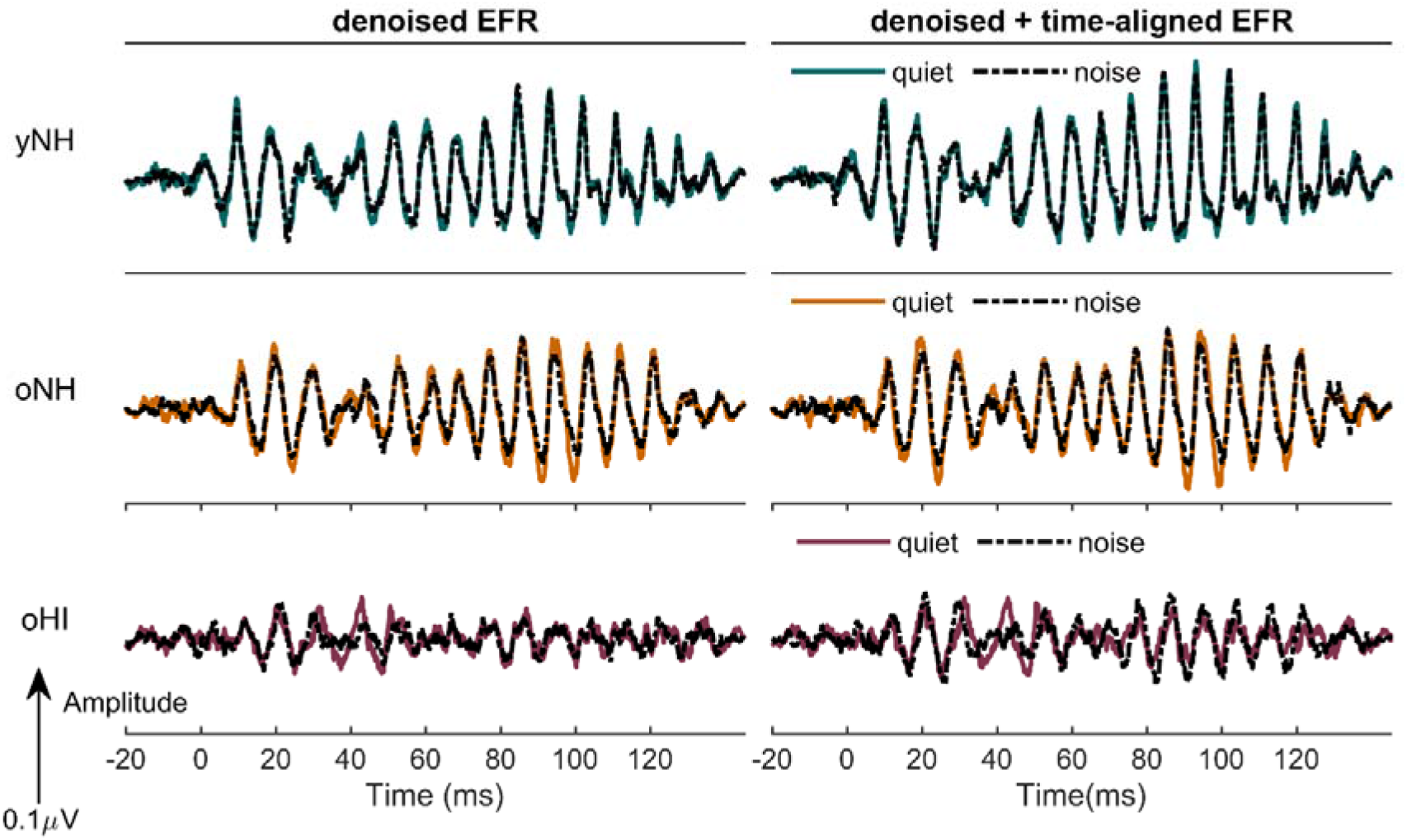
Plots of EFR pre and post bootstrapping. Grand average representations of the EFR and TFS-FFR for all three groups: younger normal hearing, older normal hearing and older hearing impaired shown on the same scales along the vertical axes and displayed from −20 ms relative to stimulus onset +20 ms relative to stimulus offset. Results from the recordings 35 dB SPL speech-weighted noise are plotted in darker lines on top of the results from the recordings in quiet.

**FIG. 6:**
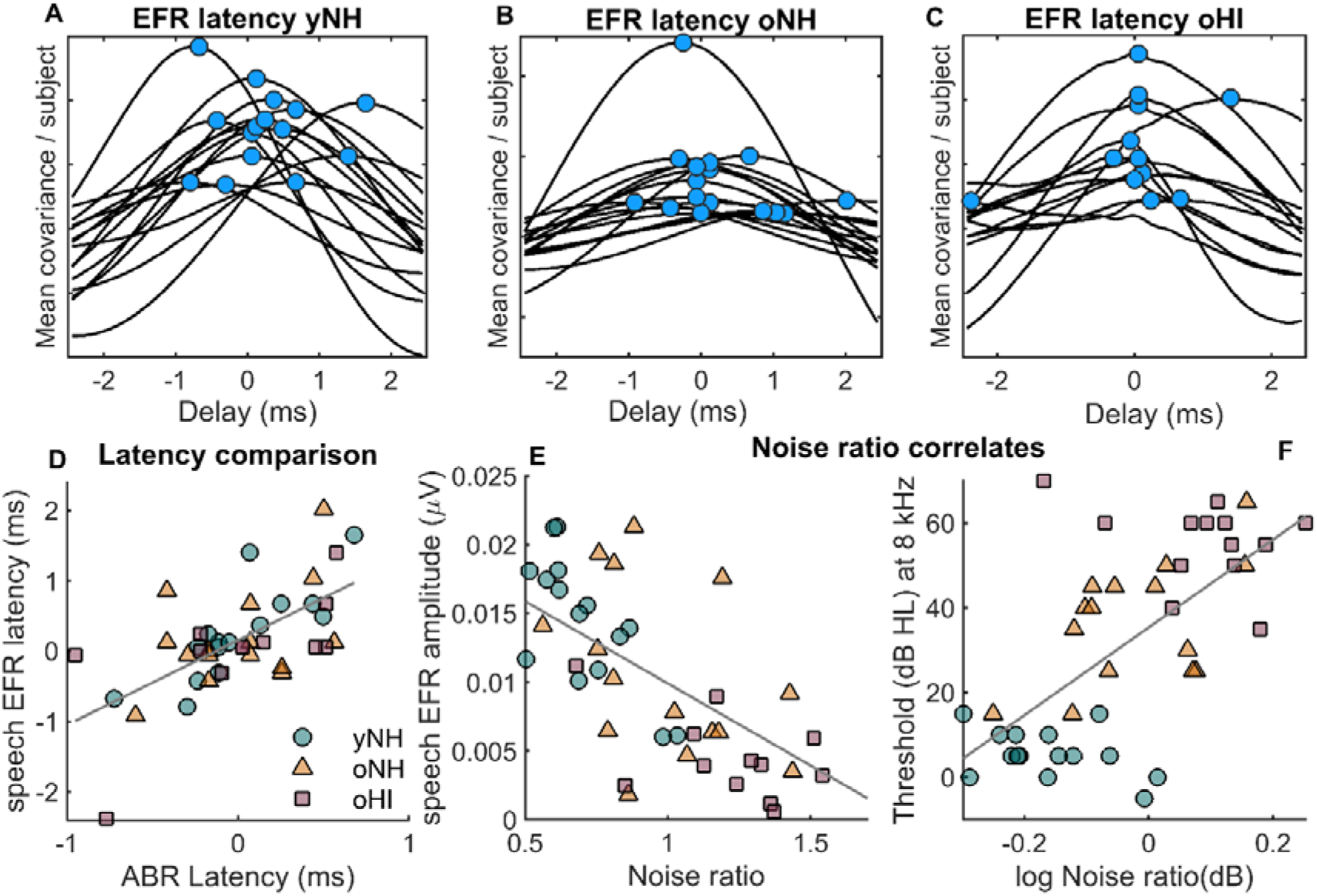
Correlation plots (II) The mean covariance as function of delay from −2.4 ms and +2.4 ms measured within each group for all subjects in A), B) and C). Blue circles denote maxima of the curves. The curves were scaled and no vertical axis was included to better visualize the curves. The yNH (square), oNH (triangle) and oHI (circle) in different colors with regression lines: D) EFR intergroup latency values as function of demeaned ABR W-V latency, E) EFR amplitude as function of noise ratio, F) Audiometric threshold at 8kHz as a function of −log(noise ratio).

The EFR and TFS-FFR averages were denoised via a bootstrapping algorithm, which was designed to preserve all of spectrotemporal information shared among opposite polarity trials (mostly envelope) or equal polarity trials (mostly fine structure). The algorithm lead to an overall improvement in reducing the noise (FIG.4), while keeping the phase of the EFR intact (FIG.5). However, the method resulted in lower values, resulting from removal of the raw spectral noise floor.

The EFR spectral magnitudes at *F*_0_ are shown in the barplots of FIG. 5. One value from one individuals from the oNH group deviated was significantly from the median values, so the data from this individual was not further used. An analysis of variance (ANOVA) confirmed that the means of the EFR magnitude at *F*_0_ were different between the three groups (F = 11.75; P <0.0001), but also when dividing the subjects in two age categories young (F = 12.85 P = 0.0009), and when dividing the groups in the categories NH and HI (F = 18.08, P = 0.0001).

To test whether the first aspect was partially caused by hidden hearing loss, and the hearing impairments dominated the difference between NH and HI, age and PTA values were used as predictors of EFR amplitude in a multiple linear regression analysis. The regression model fitted significantly to the data (R^2^ = 0.28, P = 0.0012) with an RMSE of 527 nV, and the only significant factor in the model was HF PTA (P=0.046) with a coefficient of −0.233nV/dB. Additionally, a similar trend was found when addressing the older subjects (R ^2^= 0.20, P = 0.0409) with a coefficient of −0.335nV/dB and RMSE of 522 nV, but none of the factors were able to explain the variance within the NH population.

An additional regression analysis was conducted to test whether the speech EFR was more sensitive to cochlear gain loss than the EFR responses to a rectangular wave. The speech EFR was modelled as a function of EFR_REC_, HF PTA and age in a multiple linear regression analysis. The EFR_REC_ was sufficient for modelling the speech EFR responses, but the model did not benefit from the inclusion of the other factors. The model was found significant (R^2^ = 0.55, P<0.0001) . The opposite was not true: prediction of EFR _SW_, was most optimal when including age in the model (R^2^ = 0.64, P<0.0001), with regression coefficients of 5.89 (P<0.0001) and −1.323nV/dB (P=0.0313) for the two included variables respectively.

### Latency-derived metrics

Latency was studied to detect exquisite differences between the recordings in quiet and background noise. EFR latency was derived as a within group measure by maximizing the cross covariance matrix for each individual within the group. The curves in FIG. 7 demonstrate that the specified lag window ± 2.4 ms was wide enough to find a single maximum for all subjects, while preventing the occurrence of multiple maxima. The representation of the oHI cross-covariance curves was noisier compared to the other two groups.

**FIG. 7:**
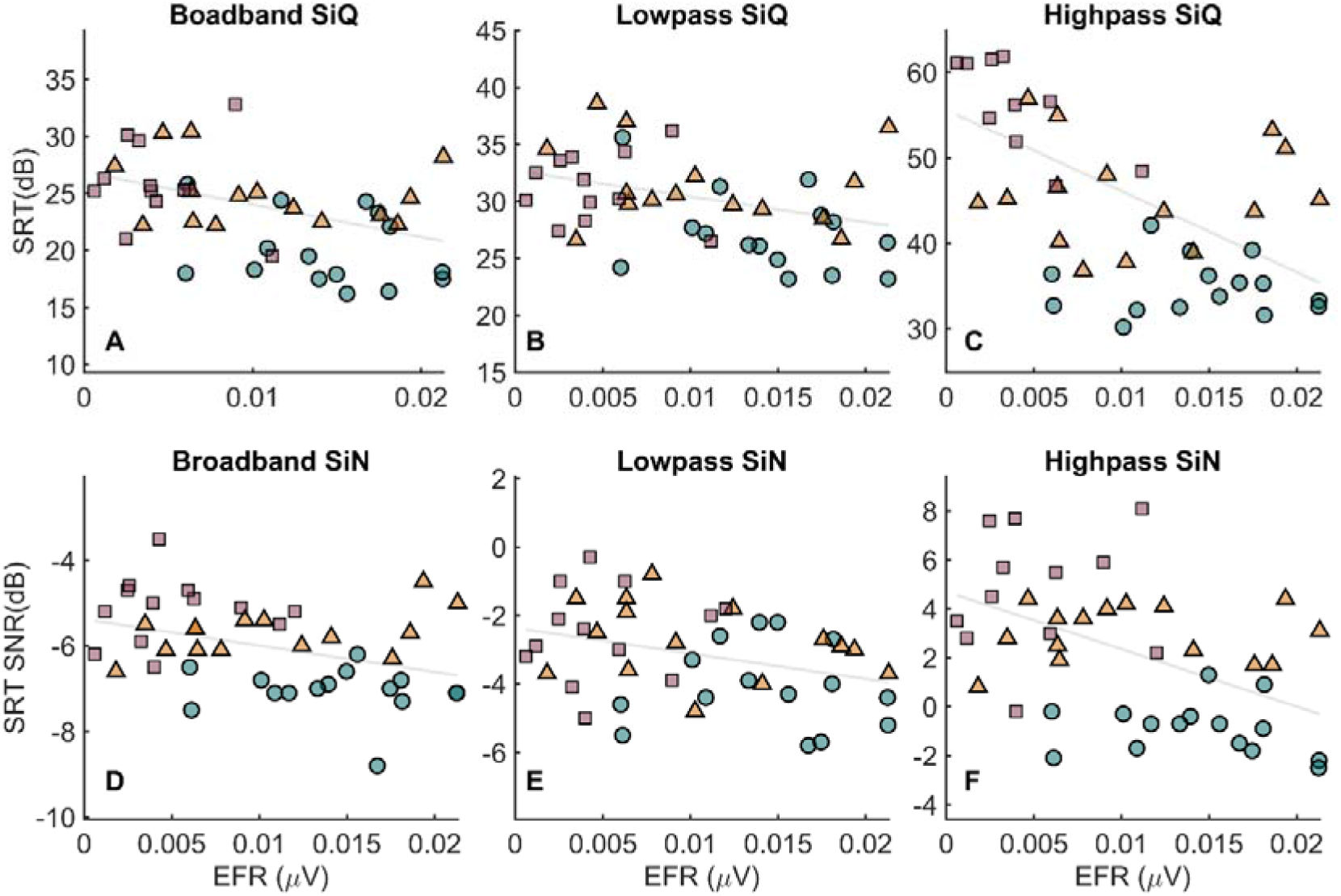
Correlation plots (II) The relation between the speech EFR (horizontal axes) and the SRT values obtained from all tested conditions for the categories yNH (square), oNH (triangle) and oHI (circle) in different colors with regression lines: A) Broadband speech with no masker, B) Lowpass filtered speech with no masker, C) Broadband speech with no masker, D) Broadband speech (80-8020 Hz) (65 dB SPL) with unfiltered SWN, E) Lowpass filtered speech with lowpass filtered SWN, F) Lowpass filtered speech with highpass filtered SWN.

Correcting for latency within the group, lead to better temporal resolution for the EFR grand averages (FIG.5). The relative improvement was determined by calculating the Hilbert transform to obtain the envelope and consequently lowpass filtering the envelope with a first order butter filter. The relative improvement was calculated as the rms of the ratio between t=0 ms and t=120 ms, resulting in improvements of 19,4%, 12.2% and 16.0% for yNH, oNH and oHI respectively and the quiet condition. Improvements were 21.3%, 14.4% and 77.5%, for the recording in noise, which was a drastic improvement for the oHI group. When comparing the final EFR grand averages, it was found via cross correlation that the EFR sustained activity was delayed with 1.3 and 1.8 ms respectively for the oNH and oHI groups compared to the yNH group.

The EFR latencies were validated with demeaned ABR W-V latency values, because the W-V peak was detected for most subjects. The best fit for modelling ABR W-V latency with EFR latency was found at the lowest intensity level of the ABR of 70 peSPL. The coefficient of determination of the fitted regression line systematically decreased as a function of stimulus level. The influence of physiological factors, such as head size, gender and age, was accounted for by running the analysis per gender and including head size diameter as a factor in the regression analysis. Adding head size resulted in a minor improvement of the regression fit and it was not significant in a multiple regression for male and female subjects for the prediction of ABR latency values, which implied that the variance in the EFR latency could mostly be attributed to subcortical delay factors (Table 1). Head size was only found significant for the yNH subjects (P=0.006) with a coefficient of 0.17 ms/ (Head diameter −54 cm).

### The effects of background noise

The effect of noise was first investigated via cross correlation. A cross correlation between the noise-free and time-aligned grand averages of the EFR, resulted in small positive latency differences of 0.12 ms and 0.24 ms for the oNH and oHI group of 0.12 ms and 0.24 ms respectively, meaning that the EFR waveform occurred later for these groups in the noise condition. There was no significant difference in magnitude between the quiet and noise condition, which was verified via paired t tests (yNH: P=0.42; oNH: P = 0.6295; oHI: P = 0.909).

As an alternative, the noise ratio was determined on the individual EFR temporal waveforms post bootstrapping and temporal alignment of each waveform. Noise ratio was expected to be a marker of noise sensitivity and therefor relate well to the SiN metrics. The index proved to be a decent marker of cochlear gain loss in NH elderly, which was demonstrated via a regression analysis with the HT at 8 kHz as predictor in FIG. 10 (R^2^ = 0.2834, P = 0.0411). This pattern was not observed for the yNH individuals, because the range of hearing thresholds at this frequency was too small. Furthermore, a linear regression analysis suggested that individuals with low noise ratios and thus a high degree of robustness to noise, were found to have generally higher EFR magnitudes. This effect was prominent in the NH population (R^2^ = 0.30, P = 0.0019), and was evident in the younger subjects (R^2^ = 0.51, P = 0.0026).

### Predictions of speech intelligibility

Predictions of SRT were formed with a linear regression analysis with the variables listed in Table 2 for the SIN conditions and shown in FIG. 8 for all conditions . The advantage of making a categorization into NH and older participants, is that the effects of ageing and cochlear gain loss could be separated.

**TABLE 2:**
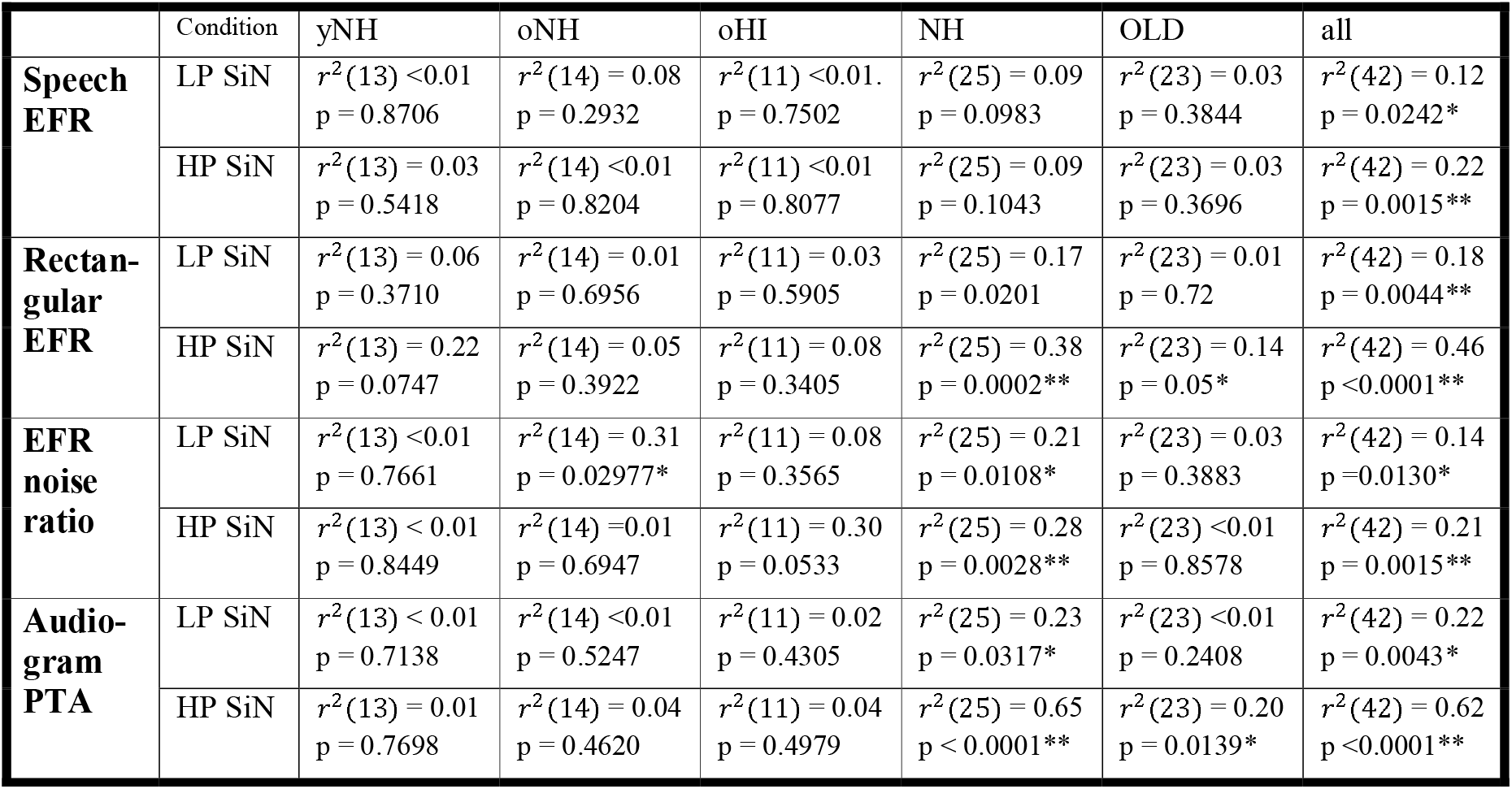
Results of speech intelligibility. Overview of the statistical results of the regression analyses, based on the regression model: SRT = X*a+b+*∈* with the data for X denoted in the first column, namely from top to bottom: speech EFR amplitude, square wave EFR amplitude, speech EFR, noise ratio and audiogram PTA values (<=1500 Hz and >1500 Hz for the LP and HP conditions respectively). And the results for Y: SRT values for the conditions are noted in the second column.* : passes the statistical tests at the significance level of 0.05 **: passes statistical confidence after Bonferroni correction for six tests (k=6) performed per condition.

Patterns for the two evaluated EFR metrics were similar: magnitude was found to be predictive of the SiN performance for all comparisons and some conditions in quiet when keeping together the entire group, but not when applying the analysis on the subgroups. These trends were present for the noise index, especially for the SRT SiN lowpass condition, but none of the relationships were statistically significant for a specific group after Bonferroni correction. Also the relation was found between SiN HP intelligibility performance and the EFR was not statistically significant for EFR_SYL_.

Consequently, the theory was tested that SRT SiN values were better explained with the audiogram than with the subcortical responses. Even though the audiogram proved to be operate on the same or higher level than the EFR measures, with equal or better prediction performances, it was still not informative enough to predict the speech intelligibility performance within each group. The audiogram was able to largely explain the notable difference in EFR magnitude between the NH and HI group, but the smaller individual differences within the groups were no good enough indicators of SRT.

### Common factors analysis

An explanatory factor analysis was performed on the EFR and speech intelligibility data. The idea was to find common factors for all parameters in terms of LF, HF hearing detection and common factors unattributable to the LF and HF PTA values, but potentially common between EFR and SRT in noise. The ten tested variables are listed in the first column of Table 3. Four factors were sufficient (P=0.093) to capture most of the variance in the dataset.

**TABLE 3:** Results of explanatory factor analysis. The explanatory factor analysis with ten independent variables and four factors. Four factors was sufficient to account for the variance in the data. The table lists all loadings and cumulative variance (communality) per variable. Loadings exceeding 0.4 are expressed in bold.

|  | *Factor 1* | *Factor 2* | *Factor 3* | *Factor 4* | *Var (%)* |
| --- | --- | --- | --- | --- | --- |
| *LF PTA* | -0.088 | **0.941** | 0.220 | 0.239 | 99.5 |
| *HF PTA* | -0.252 | **0.488** | **0.423** | **0.697** | 96.6 |
| *SRT<sub>LP</sub> SiQ* | -0.103 | **0.700** | 0.159 | 0.159 | 55.1 |
| *SRT<sub>HP</sub> SiQ* | -0.222 | **0.520** | 0.363 | **0.654** | 85.7 |
| *SRT<sub>LP</sub> SiN* | -0.198 | 0.243 | **0.629** | 0.058 | 49.7 |
| *SRT<sub>HP</sub> SiN* | -0.238 | 0.167 | **0.865** | **0.401** | 99.5 |
| *EFR<sub>syl,quiet</sub>* | **0.877** | -0.133 | -0.190 | -0.186 | 85.8 |
| *EFR<sub>syl,noise</sub>* | **0.785** | -0.074 | -0.114 | -0.039 | 63.6 |
| *EFR<sub>rec</sub>* | **0.703** | -0.119 | **-0.410** | -0.345 | 79.5 |
| *Noise ratio* | **0.465** | **-0.435** | -0.103 | **-0.489** | 65.6 |

A first observation from the table is that the first latent underlined differences in the EFR, but not in the results of the speech intelligibility tasks in noise. The second latent variable was hypothesized to be related to the ability to detect LF TFS cues near the detection threshold, while the fourth latent variable may signified the use of HF TENV cues near the detection threshold. The third latent variable was mainly responsible for differences in performance in the speech-in-noise tasks that was not shared across the EFR_SYL_ measures. The effect of noise ratio was spread towards the second and fourth latent variable, so there was likely a dependency on age related hearing deficits. However, unlike the other factors, its loading was higher than 0.4 for the first latent factor, which indicated that the EFR tended to be lower if the noise ratio was higher. Assuming that the first variable was HHL, then the results of the factor analysis would imply no link to performance in the speech in noise tasks. Instead, based on the factor loadings, it was more sensible to attribute HHL to the third latent variable, because of an inverse relation between SRT_HP_ SIN and EFR_REC_.

### Modelling supra threshold speech reception in noise

Finally, a multivariate linear model was built as a function of previously examined independent variables as predictors of SRT, focusing on the lowpass SIN and high pass SIN conditions. Variables tested in the model were the PTA values, EFR magnitude values to the speech syllable and rectangular wave and the EFR noise ratios. A first requirement of the model was to functionality explain a significant portion of the total variance in performance of speech in noise tasks. A second requirement was that the residuals between the data points and the regression line, were normally distributed around zero. The regression model was fitted to the entire data set to explain effects between the groups, as well as within each group. All possible combinations of variables as predictors were tested in the linear regression model, but the two EFR metrics, syllable and rectangular wave, were not tested together because of collinearity. Although the audiogram was capable of achieving a good performance in terms of prediction based on the regression coefficient, it lacked the property of normality, as verified with a Wilkinson-Shapiro test (P = 0.020). This was rectified by including EFR_REC_, but the overall best performance was achieved by also adding the EFR noise metric to the model. The adjusted R^2^ statistic was validated as 0.67 with a statistical confidence of P = 1.17e-10. Moreover, the hypothesis that the residuals came from a normal distribution was accepted (P = 0.31). When analyzing the SRT_LP_ SIN data set, the model of choice included the variables EFR_REC_ and LF PTA, with R^2^ of 0.23 and a statistical confidence of P = 0.0017. In addition, the residuals were normally spread around zero (P = 0.40; Kolmogorov-Smirnov). The results indicate that the responses to the speech syllable were less informative about suprathreshold speech reception than the responses to the rectangular wave.

## Discussion

### Link between the frequency following response and speech perception

This study investigated the perceptual role of the EFR and TFS-FFR in case of suprathreshold speech perception in noise for individuals from a young and older population. The older population was categorized into normal hearing and hearing impaired. The EFR to a syllable /da/ was analyzed. We found that the EFR_SYL_ was characterized by a strong reduction in magnitude in the older population and an additional stronger reduction in amplitude as a function of HF sloped hearing loss. The trends in effect size of age and hearing loss were similar in EFR_SYL_ amplitude to those found for EFR_REC,_ but there were no strong signs of an additional effect of ageing independent of cochlear gain loss.

Four metrics were tested with the aim to explain differences in speech perception performance: the EFR to a speech syllable (EFR_SYL_), a EFR noise index (NR), the EFR to a rectangular wave (EFR_REC_) and the pure tone audiogram PTA. However, EFR_REC_ proved to be a much better predictor of perceptual declines of speech perception in noise than EFR_SYL_. All variables, except, EFR_SYL_ were able to explain differences in performance of SiN highpass in the normal hearing population. This relation was important based on the known relevance of the TENV in the higher frequency domain of complex sounds (Ardoint & Lorenzi, 2010). These metrics, including PTA, shared the property that they explained differences on a whole-group level, but lacked the ability explain the variance in speech performance within each group. This finding confounded the potency of these metrics for use of individual diagnoses, instead of classification into one of the investigated groups. A factor analysis revealed that the latent variable underlining differences in EFR_SYL_ strength, was independent from the subjective audiometric results. In contrary to EFR_SYL_, the EFR shared a common latent variable with the SiN highpass condition. Combining the two variables EFR_REC_ and PTA in a multivariate linear model, resulted in the most optimal prediction for the high pass condition and explained up to 70% of the variance.

A possible reason for the decline of EFR_SYL_ in the hearing impaired group, was that the degree of cochlear synaptopathy in the oHI group was higher in respect to the oNH group. Alternatively, there is a possibility that cochlear gain loss reduced the EFR and that this behavior was not captured by the current version of the model for this type of stimulus. There is evidence that ageing degrades the ABR EFR (Fernandez et al., 2015; Keshishzadeh et al., 2020), although not every study finds an effect of age on the EFR (McClaskey et al., 2019). Additionally, recent invasive studies in mice have shown that, although ageing leads to a reduction of the sustained phase locked response at the level of spiral ganglion neurons, this effect is partially neutered by a compensatory gain mechanism effective in the IC (Parthasarathy et al., 2019). The FFR is a sum of contributions from multiple subcortical sources, including the IC, so the effects of ageing on strength of the FFR are potentially clouded by the influence of the gain mechanism. Furthermore, the reported EFR group differences based on HI and age here are much more prominent compared to the study of Goossens et al., (2019), in which an enhancement of synchronization of the steady state EFR to modulated noise maskers was reported for older hearing-impaired individuals as opposed to normal-hearing individuals, especially after controlling for hearing threshold. However, in this study, we reported a significantly smaller EFR, and a higher impact of noise in terms of latency shift and robustness to noise, which the audiogram only accounted partially for, which corroborate the findings for the rectangular-wave EFR. For future studies, it may be worthwhile to aim to separate sources from before and after the gain compensation to better study the role of the age related hidden hearing loss on the FFR. An improved separation of sources may be achieved by designing stimuli targeted to enhance the response specifically for one source or source unmixing, which could be realized by ways of source analysis or estimation of TRF (Lalor et al., 2009).

### Correlates of fine-structure and envelope in the FFR

The analysis was primarily focused on FFR magnitudes after removal of the noise floor, because magnitude was hypothesized to be more informative on cochlear synaptopathy than the SNR. There is a known dependency on the number of IHC-ANF synapses and the size of the EFR, while the impacts of age and hearing loss on the noise floor are still unclear. The TFS-FFR was not detected for most subjects in this study, but it has frequently been reported in other studies (Aiken & Picton, 2008; Mai et al., 2018). The lack of a clear TFS-FFR was substantiated by the computer model simulations. The TFS-FFR was expected to be significantly smaller in size than the EFR. The syllable was characterized by formants at lower frequency, reducing the energy at multiple harmonics of the fundamental energy. Also, the non-stable fundamental frequency lead to a more diffused representation of the energy at the harmonics, especially at higher harmonics in the power spectrum. This problem could have been circumvented by using a technique such as the Fourier analyzer (Aiken & Picton, 2008), but we specifically aimed to use the noise reduction protocol to compare the results to those of the computations in magnitude. However, it cannot be ruled out that the TFS-FFR plays a role in the detection of the syllable in for example spectral masking release tasks (Shamma & Lorenzi, 2013). Its role may be larger for speech with significant energy at higher harmonics or consonants (Léger et al., 2015).

One reason why the existence of a correlate of hidden hearing loss in the speech EFR was not proven, may be because of the nature of the EFR. According to recent research, the EFR magnitude to a vowel, is highly subject dependent, and results from cochlear interactions between basal and apical contributions (Easwar et al., 2018). The transduction of energy of a voiced sound is not perfectly synchronously as there is some delay in the excitation of the basilar membrane, which vibrates maximally at locations related to the formants. This means that auditory nerve fibers innervated at multiple locations fluctuate at the envelope modulation frequency. EEG electrodes near the scalp pick up activity from all sources, oscillating at the modulation frequency out of sync with each other. The individual differences arise from differences in inner ear morphology (e.g. length of the cochlea) or differences in the morphology of the subcortical track. The subject specific phase-sensitivity would add a substantial source of variability of EFR size, which makes it difficult to obtain a generalized measure of the present number of auditory nerve fibers A possible solution for this is averaging the EFR spectral energy density across a range of formants. We hypothesize that the predictions may be improved by averaging the responses to a subset of vowels or by using stimuli at the word or sentence level. Using a subset of vowels or words is fine in estimation of EFR strength for a proxy of the number of nerve fibers, but it will handicap estimation of the temporal waveform per vowel, as this estimation usually depends on averaging the result across thousands of repetitions. Regardless, it may provide an approach to make the EFR based predictions more accurate.

### Relevance of EFR latency study

The goal of the latency analysis was to create an enhanced view of the EFR grand averages and to study the effect of SWN. The study supported the findings that the latency of the sustained EFR matches with ABR wave V (Akhoun et al., 2008). The regression analysis also provided evidence towards the fact that this relationship is better preserved in young individuals (Parthasarathy et al., 2014), meaning that the ageing alters the representation of the FFR. The covariance matrix method was specifically developed to study the latency within a group of relatively similar individuals. However, the method was not always robust, because latency differences between the quiet and recorded sometimes exceeded 0.5 ms. The robustness of the method was linked to LF PTA, because latency differences close to zero ms always guaranteed PTA values smaller than 7dBH HL, but the differences ranged up to 2 ms for the eight out of fifteen older subjects with hearing thresholds higher than 7 dB HL.

The findings supported the claim that the subcortical sources for generation of the EFR and click ABR W-V are quite similar, most notable at lower presentation levels than at higher stimulus levels. Higher intensities lead to overall lower subcortical delays and smaller inter-subject differences. The latency study also provides evidence towards the fact that HS fibers have the largest contribution to both EFR (Encina-Llamas et al., 2019) because the contribution of the LS fibers in the click ABR is lower at high intensities. In conclusion, these results suggest that the origin of the EFR is primarily subcortical, as feedback from higher areas in the EFR would probably have resulted in additional latencies. Moreover, the correspondence and proposed method open the path of measuring EFR latency differences as a marker of cochlear synaptopathy. This is promising, because ABR latency has already been shown to correspond well to cochlear synaptopathy in case of the ABR W-V (Mehraei et al., 2016).

### Sensitivity of the EFR to noise

The noise ratio was designed to be independent of EFR strength, while being sensitive to imbalances in the representation of the EFR between different measurements. The noise ratio provided a small benefit over magnitude in terms of explaining differences in speech intelligibility results between groups, but was not strongly related to the SiN indelibility results within each group. This result may be explained by the fact that the background noise was not high enough to effectively mask the stimulus and saturate a high proportion of the HS fibers. Saturation of HS fibers with wideband noise occurs at levels much higher than 35 dB rms SPL (Furman et al., 2013). Running an experiment with the signal embedded in noise is challenging, because the FFR is typically not be retrieved when the masker level approaches the signal level. Because of this limitation, positive values in the range of 5-10 dB SNR are most often preferred to study the effects of noise (e.g. Bidelman, 2016). However, even despite the low noise level, the trends in the table were overall similar when comparing the results obtained for the audiogram and EFR_REC._ The factor analysis proved that the latent factors were shared among the first two latent factors, which indicated that the noise analysis was potentially informative on the ability to effectively use LF cues and HF cues.

## List of abbreviations

ABR: auditory brainstem response
ANF: auditory nerve fiber
CS: cochlear synaptopathy
EFR: envelope following response
FFR: frequency following response
HF: high frequency
HI: hearing-impaired
HHL: hidden hearing loss
HS: high spontaneous
LF: low frequency
LS: low spontaneous
MS: medium spontaneous
NH: normal-hearing
OHC: outer hair cell loss
peSPL: peak-equivalent sound pressure level
PTA: pure tone average
SNHL: sensorineural hearing loss
SRT: speech reception threshold
SiN: speech in noise
SiQ: speech in quiet
TENV: temporal envelope
TFS: temporal fine structure

## Summary

Bundling all of the findings of this study, the results can be summarized as follows:

- **Hearing impairment** The older subjects were characterized by a lower EFR_SYL._ Also HF sloped hearing loss reduced the EFR, but this difference was not necessarily explained by HF PTA. The effect of noise on the EFR was also higher in the older categories, in terms of delay and robust encoding.
- **Speech perception in noise** The group EFR averages were found to be proportional to group SiN performance .The EFR was maximal for yNH, followed by oNH and oHI. The EFR_REC_ was less sensitive to audiogram than EFR_SYL._ EFR_REC_ was found to be mostly related to the SRT_HP_ SiN, while EFR_SYL_ to the HF PTA, SRT_HP_ SiN and SRT_LP_ SiN.
- **Latency** Obtained EFR latency values, derived from cross covariance computation, correlated well with click ABR W-V latency values. The derived latency values were also useful for improved visualization purposes of the grand averages of the EFR. Moreover, we found a latency shift in the EFR in response to background noise for the older subjects.

---

**Supplementary.**
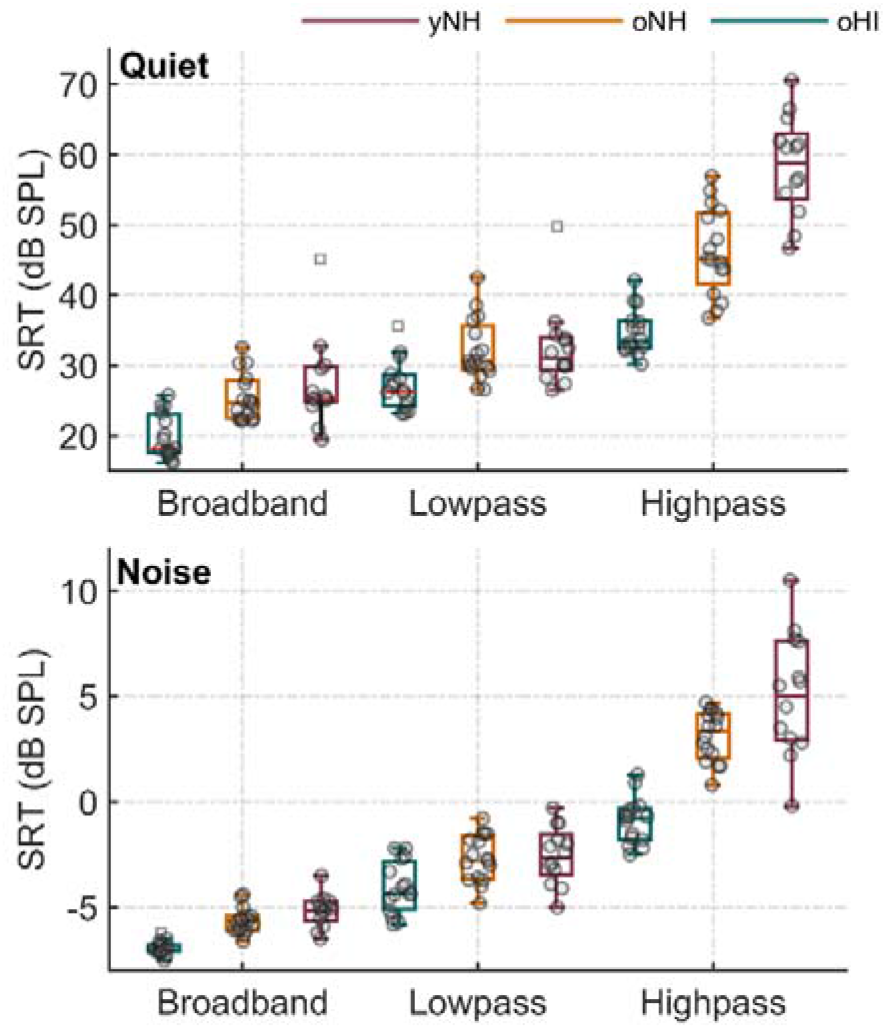
Speech in Noise performances (barplot) The results of the speech intelligibility tests in boxplots with medians, quartiles and whiskers with the interquartile ranges. Boxes indicate SRT values for all the tests performed in quiet and in stationary background noise for different filters applied to the sentences and to the background noise.

## References

Aiken, S. J., & Picton, T. W. (2008). Envelope and spectral frequency-following responses to vowel sounds. Hearing Research, 245(1–2), 35–47. https://doi.org/10.1016/j.heares.2008.08.004

Akhoun, I., Gallégo, S., Moulin, A., Ménard, M., Veuillet, E., Berger-Vachon, C., Collet, L., & Thai-Van, H. (2008). The temporal relationship between speech auditory brainstem responses and the acoustic pattern of the phoneme /ba/ in normal-hearing adults. Clinical Neurophysiology. https://doi.org/10.1016/j.clinph.2007.12.010

Anderson, S., Skoe, E., Chandrasekaran, B., & Kraus, N. (2010). Neural Timing Is Linked to Speech Perception in Noise. Journal of Neuroscience, 30(14), 4922–4926. https://doi.org/10.1523/JNEUROSCI.0107-10.2010

Ardoint, M., & Lorenzi, C. (2010). Effects of lowpass and highpass filtering on the intelligibility of speech based on temporal fine structure or envelope cues. Hearing Research, 260(1–2), 89–95. https://doi.org/10.1016/j.heares.2009.12.002

Bharadwaj, H. M., Masud, S., Mehraei, G., Verhulst, S., & Shinn-Cunningham, B. G. (2015). Individual differences reveal correlates of hidden hearing deficits. Journal of Neuroscience. https://doi.org/10.1523/JNEUROSCI.3915-14.2015

Bharadwaj, H. M., Verhulst, S., Shaheen, L., Charles Liberman, M., & Shinn-Cunningham, B. G. (2014). Cochlear neuropathy and the coding of supra-threshold sound. Frontiers in Systems Neuroscience. https://doi.org/10.3389/fnsys.2014.00026

Bidelman, G. M. (2016). Relative contribution of envelope and fine structure to the subcortical encoding of noise-degraded speech. The Journal of the Acoustical Society of America, 140(4), EL358–EL363. https://doi.org/10.1121/1.4965248

Carney, L. H. (2018). Supra-Threshold Hearing and Fluctuation Profiles: Implications for Sensorineural and Hidden Hearing Loss. In JARO - Journal of the Association for Research in Otolaryngology. https://doi.org/10.1007/s10162-018-0669-5

Choi, J. M., Purcell, D. W., Coyne, J. A. M., & Aiken, S. J. (2013). v. Ear and Hearing, 34(5), 637–650. https://doi.org/10.1097/AUD.0b013e31828e4dad

Coffey, E. B. J., Colagrosso, E. M. G., Lehmann, A., Schönwiesner, M., & Zatorre, R. J. (2016). Individual Differences in the Frequency-Following Response: Relation to Pitch Perception. PloS One. https://doi.org/10.1371/journal.pone.0152374

Easwar, V., Banyard, A., Aiken, S., & Purcell, D. (2018). Phase delays between tone pairs reveal interactions in scalp-recorded envelope following responses. Neuroscience Letters. https://doi.org/10.1016/j.neulet.2017.12.014

Easwar, V., Purcell, D. W., Aiken, S. J., Parsa, V., & Scollie, S. D. (2015). Evaluation of Speech-Evoked Envelope Following Responses as an Objective Aided Outcome Measure: Effect of Stimulus Level, Bandwidth, and Amplification in Adults with Hearing Loss. Ear and Hearing. https://doi.org/10.1097/AUD.0000000000000199

Encina-Llamas, G., Harte, J. M., Dau, T., Shinn-Cunningham, B., & Epp, B. (2019). Investigating the Effect of Cochlear Synaptopathy on Envelope Following Responses Using a Model of the Auditory Nerve. JARO - Journal of the Association for Research in Otolaryngology. https://doi.org/10.1007/s10162-019-00721-7

Fernandez, K. A., Jeffers, P. W. C., Lall, K., Liberman, M. C., & Kujawa, S. G. (2015). Aging after Noise Exposure: Acceleration of Cochlear Synaptopathy in “Recovered” Ears. Journal of Neuroscience, 35(19), 7509–7520. https://doi.org/10.1523/JNEUROSCI.5138-14.2015

Furman, A. C., Kujawa, S. G., & Liberman, M. C. (2013). Noise-induced cochlear neuropathy is selective for fibers with low spontaneous rates. Journal of Neurophysiology, 110(3), 577–586. https://doi.org/10.1152/jn.00164.2013

Goossens, T., Vercammen, C., Wouters, J., & van Wieringen, A. (2019). The association between hearing impairment and neural envelope encoding at different ages. Neurobiology of Aging. https://doi.org/10.1016/j.neurobiolaging.2018.10.008

Henry, K. S., Kale, S., & Heinz, M. G. (2014). Noise-induced hearing loss increases the temporal precision of complex envelope coding by auditory-nerve fibers. Frontiers in Systems Neuroscience, 8(20), 1–10. https://doi.org/10.3389/fnsys.2014.00020

Joris, P. X., & Verschooten, E. (2013). On the limit of neural phase locking to fine structure in humans. Advances in Experimental Medicine and Biology. https://doi.org/10.1007/978-1-4614-1590-9-12

Keshishzadeh, S., Garrett, M., Vasilkov, V., & Verhulst, S. (2020). The Derived-Band Envelope Following Response and its Sensitivity to Sensorineural Hearing Deficits. BioRxiv. https://doi.org/10.1101/820704

Kollmeier, B., Warzybok, A., Hochmuth, S., Zokoll, M. A., Uslar, V., Brand, T., & Wagener, K. C. (2015). The multilingual matrix test: Principles, applications, and comparison across languages: A review. In International Journal of Audiology. https://doi.org/10.3109/14992027.2015.1020971

Lalor, E. C., Power, A. J., Reilly, R. B., & Foxe, J. J. (2009). Resolving Precise Temporal Processing Properties of the Auditory System Using Continuous Stimuli. Journal of Neurophysiology. https://doi.org/10.1152/jn.90896.2008

Léger, A. C., Reed, C. M., Desloge, J. G., Swaminathan, J., & Braida, L. D. (2015). Consonant identification in noise using Hilbert-transform temporal fine-structure speech and recovered-envelope speech for listeners with normal and impaired hearing. The Journal of the Acoustical Society of America, 138(1), 389–403. https://doi.org/10.1121/1.4922949

Mai, G., Tuomainen, J., & Howell, P. (2018). Relationship between speech-evoked neural responses and perception of speech in noise in older adults. The Journal of the Acoustical Society of America. https://doi.org/10.1121/1.5024340

McClaskey, C. M., Dias, J. W., & Harris, K. C. (2019). Sustained envelope periodicity representations are associated with speech-in-noise performance in difficult listening conditions for younger and older adults. Journal of Neurophysiology. https://doi.org/10.1152/jn.00845.2018

Mehraei, G., Hickox, A. E., Bharadwaj, H. M., Goldberg, H., Verhulst, S., Liberman, M. C., Barbara, X., & Shinn-Cunningham, G. (2016). Auditory brainstem response latency in noise as a marker of cochlear synaptopathy. The Journal of Neuroscience, 36(13), 3755–3764. https://doi.org/10.1523/JNEUROSCI.4460-15.2016

Mepani, A. M., Verhulst, S., Hancock, K. E., Garrett, M., Vasilkov, V., Bennett, K., de Gruttola, V., Liberman, M. C., & Maison, S. F. (2021). Envelope following responses predict speech-in-noise performance in normal-hearing listeners. Journal of Neurophysiology, 125(4). https://doi.org/10.1152/jn.00620.2020

Parthasarathy, A., Datta, J., Torres, J. A. L., Hopkins, C., & Bartlett, E. L. (2014). Age-related changes in the relationship between auditory brainstem responses and envelope-following responses. JARO - Journal of the Association for Research in Otolaryngology. https://doi.org/10.1007/s10162-014-0460-1

Parthasarathy, A., Herrmann, B., & Bartlett, E. L. (2019). Aging alters envelope representations of speech-like sounds in the inferior colliculus. Neurobiology of Aging. https://doi.org/10.1016/j.neurobiolaging.2018.08.023

Paul, B. T., Bruce, I. C., & Roberts, L. E. (2016). Evidence that hidden hearing loss underlies amplitude modulation encoding deficits in individuals with and without tinnitus. Hearing Research. https://doi.org/10.1016/j.heares.2016.11.010

Ruggles, D., Bharadwaj, H., & Shinn-Cunningham, B. G. (2012). Why middle-aged listeners have trouble hearing in everyday settings. Current Biology, 22(15), 1417–1422. https://doi.org/10.1016/j.cub.2012.05.025

Schoof, T., & Rosen, S. (2016). The Role of Age-Related Declines in Subcortical Auditory Processing in Speech Perception in Noise. Journal of the Association for Research in Otolaryngology, 17(5), 441–460. https://doi.org/10.1007/s10162-016-0564-x

Shamma, S., & Lorenzi, C. (2013). On the balance of envelope and temporal fine structure in the encoding of speech in the early auditory system. The Journal of the Acoustical Society of America, 133(5), 2818–2833. https://doi.org/10.1121/1.4795783

Vasilkov, V., Garrett, M., Mauermann, M., & Verhulst, S. (2021). Enhancing the sensitivity of the envelope-following response for cochlear synaptopathy screening in humans: The role of stimulus envelope. Hearing Research, 400. https://doi.org/10.1016/j.heares.2020.108132

Verhulst, S., Altoè, A., & Vasilkov, V. (2018). Computational modeling of the human auditory periphery: Auditory-nerve responses, evoked potentials and hearing loss. In Hearing Research. https://doi.org/10.1016/j.heares.2017.12.018

Verhulst, S., Jagadeesh, A., Mauermann, M., & Ernst, F. (2016). Individual Differences in Auditory Brainstem Response Wave Characteristics: Relations to Different Aspects of Peripheral Hearing Loss. Trends in Hearing. https://doi.org/10.1177/2331216516672186

Verhulst, S., & Warzybok, A. (2018). Contributions of Low- and High-Frequency Sensorineural Hearing Deficits to Speech Intelligibility in Noise. BioRxiv.

Wagener, K., Brand, T., & Kollmeier, B. (1999). Entwicklung und Evaluation eines Satztests für die deutsche Sprache Teil III: Optimierung des Oldenburger Satztests. Z Audiol. https://doi.org/10.1039/C4LC00330F

Wardenga, N., Batsoulis, C., Wagener, K. C., Brand, T., Lenarz, T., & Maier, H. (2015). Do you hear the noise? the German matrix sentence test with a fixed noise level in subjects with normal hearing and hearing impairment. International Journal of Audiology. https://doi.org/10.3109/14992027.2015.1079929

Young, E. D., & Sachs, M. B. (1979). Representation of steady-state vowels in the temporal aspects of the discharge patterns of populations of auditory-nerve fibers. The Journal of the Acoustical Society of America, 66(5), 1381–1403. https://doi.org/10.1121/1.383532

Zhu, L., Bharadwaj, H., Xia, J., & Shinn-Cunningham, B. (2013). A comparison of spectral magnitude and phase-locking value analyses of the frequency-following response to complex tones. The Journal of the Acoustical Society of America. https://doi.org/10.1121/1.4807498

